# Two Novel Forms of ERG Oscillation in *Drosophila*: Age and Activity Dependence

**DOI:** 10.1101/259952

**Authors:** Atsushi Ueda, Scott Woods, Ian McElree, Tristan C.D.G. O’Harrow, Casey Inman, Savantha Thenuwara, Muhammad Aftab, Atulya Iyengar

## Abstract

Over an animal’s lifespan, neuronal circuits and systems often decline in an inherently heterogeneous fashion. To compare the age-dependent progression of changes in visual behavior with alterations in retinal physiology, we examined phototaxis and electroretinograms (ERGs) in a wild-type *D. melanogaster* strain *(Canton-S)* across their lifespan. In aged flies (beyond 50% median lifespan), we found a marked decline in phototaxis, while motor coordination was less disrupted, as indicated by relatively stronger negative geotaxis. These aged flies displayed substantially reduced ERG transient amplitudes while the receptor potentials (RP) remained largely intact. Using a repetitive light flash protocol, we serendipitously discovered two forms of activity-dependent oscillation in the ERG waveforms of young flies: “light-off’ and “light-on” oscillations. After repeated 500 ms light flashes, light-off oscillations appeared during the ERG off-transients (frequency: 50-120 Hz, amplitude: ~1 mV). Light-on oscillations (100-200 Hz, ~0.3 mV) were induced by a series of 50 ms flashes, and were evident during the ERG on-transients. Both forms of oscillation were observed in other strains of *D. melanogaster(Oregon-R, Berlin)*, additional *Drosophila* species *(funerbris, euronotus, hydei, americana)*, and were evoked by a variety of light sources. Both light-off and light-on oscillations were distinct from previously described ERG oscillations in visual mutants, such as *rosA*, in terms of location within the waveform and frequency. However, within *rosA* mutants, light-off oscillations, but not light-on oscillations could be recruited by the repetitive light flash protocol. Importantly though, we found that both forms of oscillation were rarely observed in aged flies. Although the physiological bases of these oscillations remain to be elucidated, they may provide important clues to age-related changes in neuronal excitability and synaptic transmission.

## Introduction

The *Drosophila* electroretinogram (ERG) provides an accessible and incisive readout of visual system function in a genetically tractable model organism (Pak, 1975; Stark & Wasserman, 1974; Vilinsky & Johnson, 2012). The complex waveform represents an extracellular combination of electrical activities associated with phototransduction and synaptic transmission. Prominent features of the ERG waveform include a sustained receptor potential (RP), as well as light on- and light off-transients, with each feature corresponding to a distinct set of underlying physiological processes. The RP component represents the sustained depolarization in photoreceptor cells (Heisenberg, 1971; Alawi & Pak, 1971), while the transients generally reflect synaptic potentials of the photoreceptor cells and their targets, large monopolar neurons in the optic lamina (Heisenberg, 1971; Coombe, 1986). Recent studies on neurotransmitter re-uptake by adjacent glia have also implicated these cells in shaping the ERG waveform (Rahman, *et al*., 2012; Chaturvedi, Reddig, & Li, 2014). Thus, features of the *Drosophila* ERG waveform serve as a report on the performance of photoreceptors, 2^nd^ order neurons, and surrounding glia in initial visual information processing.

Mutant lines with defective ERG waveforms have provided significant insight into the functions of a number of genes which encode components of 2^nd^ messenger systems, ion channels, and synaptic transmission machinery (Pak, 1975; Pak, 2010). Notable examples include mutants of *no receptor potential A (norpA)*, encoding Phospholipase C (Bloomquist *et al*., 1988); and *transient receptor potential* (*trp*, Minke, Wu, & Pak, 1975), encoding the founding member of the Na^+^/Ca^2^+-permeable TRP superfamily of cation channels (Montell & Rubin, 1989; Clapham, Runnels, & Strubing, 2001). With the advent of transgenic techniques, *Drosophila* ERGs have proven themselves invaluable by providing first-order assessments of neuronal dysfunction due to disease-associated mutations (Yamamoto *et al*., 2014; Oortveld *et al*, 2013). Indeed, overexpression of mutant proteins including α-Synuclein, (Chouhan *et al*., 2016), Parkin (West, Elliott, & Wade, 2015), and Huntingtin (Lee, Yoshihara, & Littleton, 2003) often leads to age-dependent disruptions of synaptic transmission and cellular morphology, evident in altered ERG waveforms. Despite the widespread use of the ERG in studying the genetics of age-related neurodegeneration, the progression of alterations that the waveform undergoes during normal healthy aging remains to be fully characterized.

As part of the course *Neurobiology Laboratory* at the University of Iowa, we performed ERGs on wild-type (WT) and several mutant *Drosophila* strains across their lifespan. Groups of two or three students each designed a series of experiments to assess the impact of genetic perturbations on age-related changes in visual behavior and physiology against WT and relevant controls. We utilized Benzer’s countercurrent apparatus (Benzer, 1967) to assess phototaxis, and we performed ERGs to uncover changes in retinal physiology. The technical simplicity of ERG recordings coupled with the physiological insight they provide make the approach ideal for introducing basic concepts of electrophysiology. Due to time constraints and limited sample sizes, individual student groups were oftentimes not able to develop robust conclusions based on their data alone. However, by pooling results from common genotypes across student groups and by conducting a series of follow-up experiments, we were able to compare the timing of age-related changes in visual behavior with changes in ERG properties. Furthermore, using a repetitive light stimulus protocol consisting of trains of long-duration (500 ms) or short-duration (50 ms) flashes with varying inter-flash intervals (0.1 - 2.5 s), we discovered two novel forms of oscillation in the *Drosophila* ERG signal. Together, our results highlight the potential for discovering novel age-dependent phenomena in the *Drosophila* ERG waveform.

## Materials and methods

### Drosophila husbandry

Flies were collected 0-3 days post-eclosion, and were housed in standard vials containing cornmeal-agar medium (Frankel & Brosseau, 1968), replaced at least once a week. To accelerate age-related changes in physiology, the initial cohorts of WT flies were reared within a 29 °C incubator, under constant darkness (apart from door openings and during transferring). All other flies were reared at 23 °C under a 12:12 hr light-dark conditions. The WT strain used for longevity experiments was *Canton-S* (Ruan & Wu, 2008). Other strains used include the wild-type strains *Oregon-R*, and *Berlin*; as well as the mutant strains *rosA*^*p*213^; Burg, Geng, Guan, Koliantz & Pak 1996). The populations of *Drosophila* species, *D. funebris, D. euronotus* and *D. americana*, were gifts from Dr. Bryant McAllister. The *D. hydei* strain was acquired from Dr. Chun-Fang Wu’s stock collection. All strains used had wild-type eye pigmentation.

### Phototaxis & negative geotaxis

The countercurrent apparatus used for negative geotaxis and phototaxis assays was originally developed by Seymour Benzer (1967). A dim red light in the room facilitated machine operation. Between 7 and 40 flies were loaded into the starting tube of a four-tube countercurrent machine. For phototaxis assays, the apparatus was placed horizontally in a light box, with an LED strip light (SuperNight 5050 LEDs, Ebestrade, Portland, OR) placed approximately 2 cm from the machine. For negative geotaxis assays, the apparatus was positioned vertically. To start each round of taxis, the machine was “banged down” to settle the flies. The flies were allowed to move to the opposite tube for 20 s, after which the opposite tube was advanced to the next tube of the apparatus. After three rounds, the number of flies able to successfully traverse to the opposite tube zero, one, two or three times was recorded. The phototaxis and negative geotaxis indices were computed by finding the average number of tubes traversed by flies of a given age.

### ERG recordings

ERG recording procedures have been described previously (Pak, Grossfield, & White, 1969; Dolph, Nair, & Raghu, 2010). Anesthetized flies were mounted into the holes of a breadboard to restrain movement and expose the head. Melted polyethylene glycol wax (melting point: 45-50 °C) was applied to the back of the head to secure the fly to the breadboard. Initial experiments utilized ethyl ether as an anesthetic while later experiments used CO_2_. We did not detect any differences in ERG waveforms based on which anesthetic was used. Flies were allowed to recover at least 15 min prior to ERG recordings.

Electrodes were constructed from filamented glass micropipettes (1.22 mm OD, 0.68 mm ID, WPI, Sarasota, FL) using an electrode-puller (Model PP-83, Narishige Scientific Instrument Lab, Tokyo, Japan). Electrodes were filled with a saline solution (0.7% w/v NaCl) and inserted into an electrode holder containing a chloridized silver wire. The electrode resistance was ~1 MO. The recording electrode was inserted into the cornea at an approximately normal angle. The ground electrode tip was broken and inserted into the proboscis. Signals were amplified 10x by a DC amplifier (IX1, Dagan Corporation, Minneapolis, USA). The amplified signal was sampled at 10 kHz by a data acquisition device (PowerLab 26T, ADInstruments, Colorado Springs CO, USA) connected to a PC running LabChart software (ADInstruments).

Unless otherwise noted, light flash trains were delivered by a 5 mm generic cool-white LED driven at constant voltage. The LED was positioned towards the eye, approximately 4 cm away, and the light intensity at the eye was ~1,200 lux. Every fly examined underwent six light flash trains, each train consisting of 60 s of dark adaptation followed by ten light flashes. Light flashes during each train were either 50 or 500 ms long, and had an inter-flash interval of 0.1, 0.5 or 2.5 s.

### Data analysis

ERG traces were analyzed using custom-written MATLAB scripts (Mathworks, Natick, MA) to quantify on-transients, off-transients, receptor potentials and other notable features.

## Results

### Age-related changes in phototaxis and geotaxis

Progressive decline in behavioral performance is commonly associated with aging. Using Benzer’s countercurrent apparatus (1967), we examined the phototactic responses of two populations of WT *(Canton-S)* flies with distinct lifespan curves: A high temperature-reared group (29 °C) with a compressed lifespan (median ~30 d), and a room temperature-reared group (23 °C) with a relatively longer lifespan (median ~65 d). At both rearing temperatures, we observed initially strong phototactic response in young flies followed by a progressive decline in phototactic behavior across the lifespan (Figure 1). Indeed, beyond 50% of the median lifespan for the two rearing temperatures (15 d and 32 d respectively), the phototaxis index was substantially reduced compared to younger counterparts. To determine if the observed decline in phototactic response was due to a corresponding loss of motor coordination, we subjected the same populations of flies to a negative geotaxis assay using the same countercurrent apparatus (see Methods). Importantly, across ages displaying marked decline in phototaxis, the negative geotaxis indices were consistently higher than corresponding phototaxis indices. This difference was most prominent at 18 d for 29 °C-reared flies (1.4 vs 2.2, Figure 1A) and at 30 d for 23 °C-reared flies (0.4 vs 2.0, Figure 1B). Our findings suggests that age-related decline in phototaxis precedes general loss of motor coordination, and may be due to alterations in visual system function.

**Figure 1.**
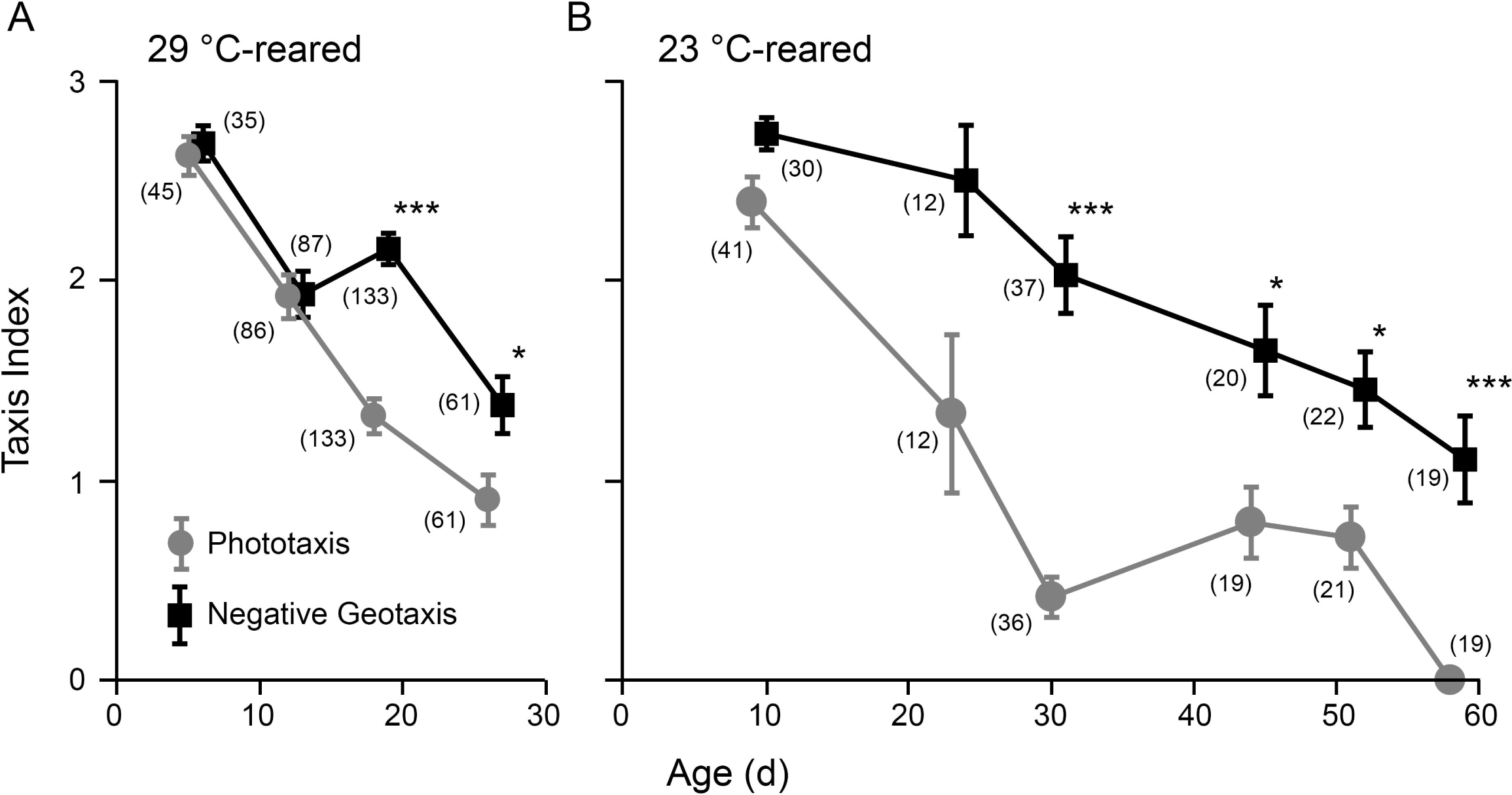
Distinct progression of age-dependent decline in phototaxis and negative geotaxis. (**A**) 29 °C-reared individuals. (**B**) 23 °C-reared individuals. Points represent the mean photo/geo-taxis index for WT flies. Error bars indicate SEM. (* *p* < 0.05, *** *p* < 0.001, Kruskal-Wallis ANOVA, Bonferroni-corrected rank-sum *post hoc* test. n = 7 - 40 flies per run, 2 - 4 runs per age, number of total flies per age group as indicated in parenthesis)

### Age-related changes in ERG

To correlate the observed decline in visual system function with changes in retinal physiology, we recorded ERGs from 29 °C-reared WT flies across their lifespan (Figure 2). In young flies responding to a 500 ms light flash, we observed robust receptor potentials (RP, ~10 mV) as well as on-transients (~2 mV) and off-transients (~ 5 mV, Figure 2A). Across all ages examined, we found the RP amplitude was largely maintained (Figure 2B). However, as flies aged, we observed a substantial reduction in the amplitudes of on- and off-transients (Figure 2A). Indeed, in most flies older than 15 d, the on-transients were undetectable (5/8 flies, Figure 2C) and the off-transients were less than 1 mV (7/8 flies, Figure 2D) in amplitude. Notably, additional light flashes did not evoke transients in older flies initially lacking transients. To extend our analysis of age-related changes in ERG waveforms, we also recorded ERGs across the lifespan of the longer-lived 23 °C-reared population (Supplemental Figure 1). Similar to the observed trends in the 29 °C-reared population, we found that the receptor potential amplitude was largely stable across the lifespan, and also noted a reduction in on- and off-transient amplitudes in the oldest flies compared to their younger counterparts. However, in most cases, both on- and off-transients still were detectable in aged individuals, consistent with previous observations (Phillips, Woodruff, Liang, Pattern, & Broadie, 2008; Jaiswal *et al*., 2015). These findings suggest that over the course of normal-healthy aging, synaptic transmission between photoreceptors and 2^nd^ order targets in the lamina are selectively disrupted prior to any decline in the phototransduction cascade leading to the receptor potential.

**Figure 2.**
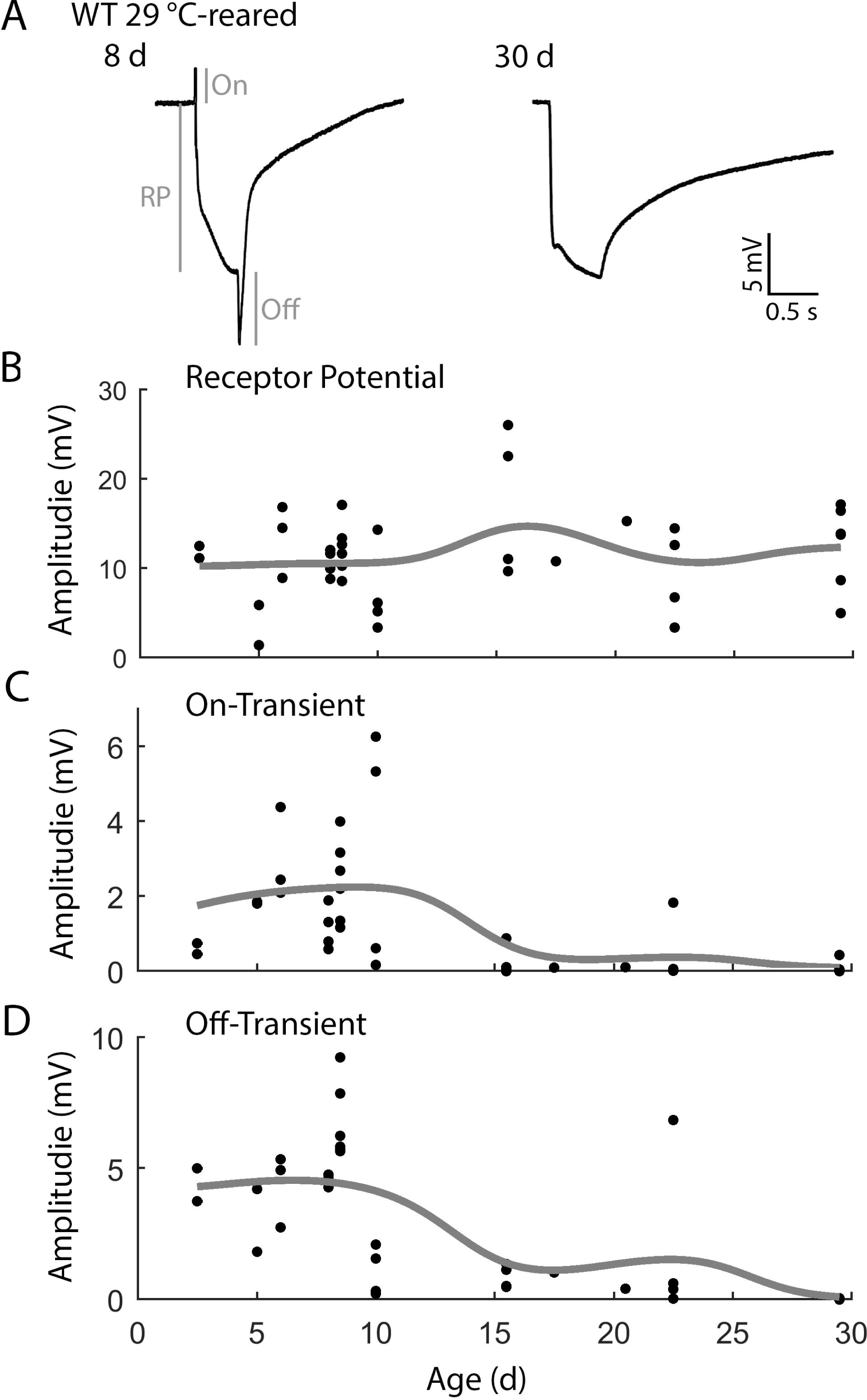
Age-dependent alterations of ERG waveforms in 29 °C-reared WT flies. (**A**) Representative ERGs from 8 and 30 d flies triggered by a 500 ms light flash. ERG characteristics analyzed include receptor potential (RP), on-transient (On), and off-transient (Off) as depicted in the 8 d sample ERG. Note that on- and off-transients are attenuated in the 30 d example. (**B-D**) Age-dependent alterations in the amplitudes of (**B**) RP, (**C**) on-transient, and (**D**) off-transients. Trend-lines indicate a Gaussian-kernel running average (σ = 5 d). n = 42 flies.

### Two novel forms of ERG waveform oscillation

During the course of our ERG experiments, we discovered that repetitive light flash protocols (flash duration: 50, or 500 ms; inter-flash interval: 2.5, 0.5 or 0.1 s; Figure 3A) led to two previously undocumented forms of oscillation in the ERG waveforms of young WT flies. The first form of oscillation appeared during ERG responses to 500-ms flashes. Although the initial ERG waveforms generally appeared to be normal, successive waveforms often displayed a distinctive oscillation during the off-transient (Figure 3B). These “light-off oscillations” had a frequency between 70 and 110 Hz and an amplitude that appeared to grow upon successive stimulations, suggesting an activity-dependent recruitment. Indeed, we found that these oscillations were most obvious, up to 1 mV in amplitude, between the 8^th^ and 10^th^ light flash. Upon closer examination, however, several cases of subtle light-off oscillation were found in the off-transient of the first flash as well.

**Figure 3.**
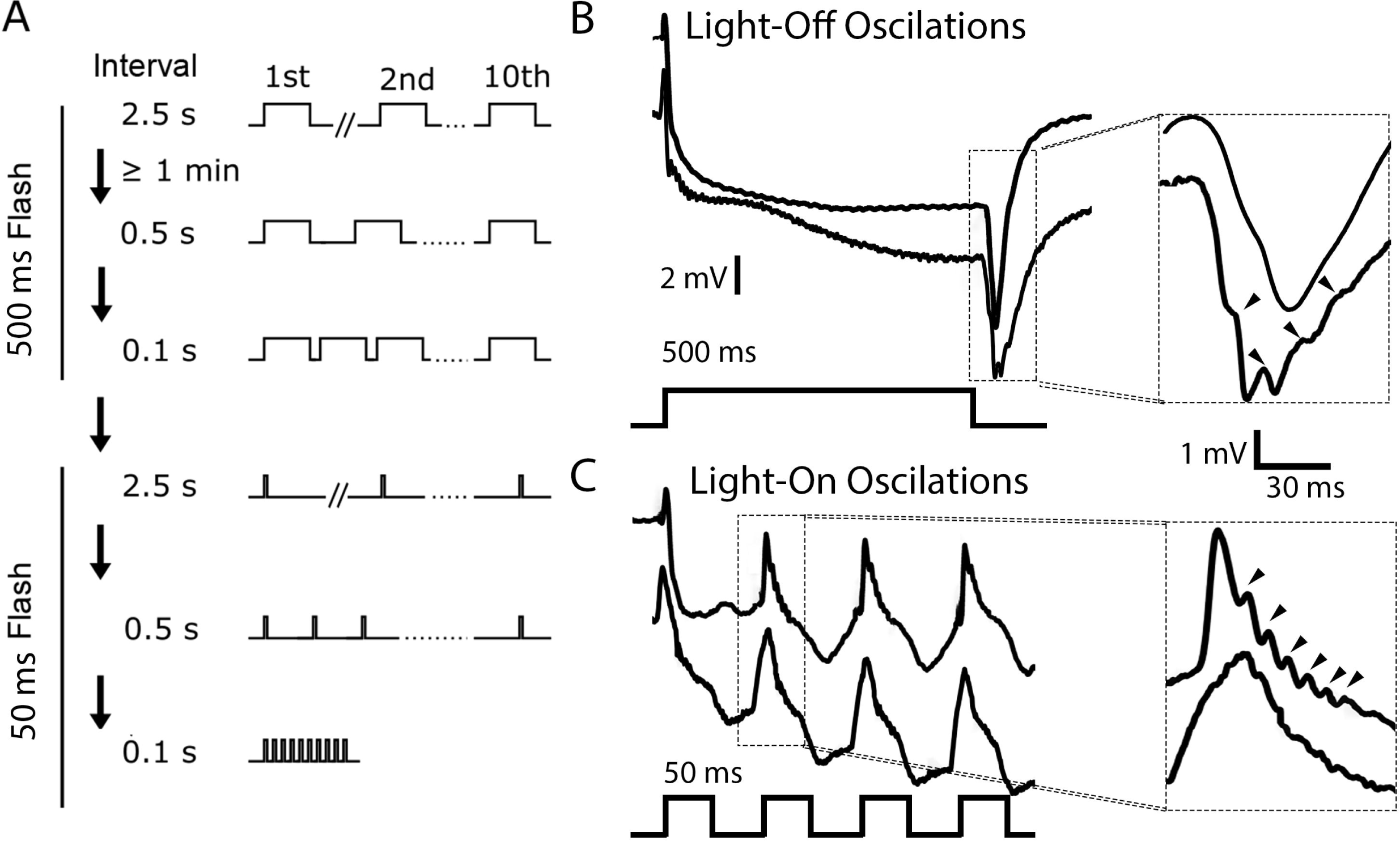
ERG oscillations triggered by repetitive light flashes. (**A**) Repetitive light flash protocol. Six 10-flash stimulus trains were applied to each fly. The first set of three trains used 500 ms flashes delivered at 2.5, 0.5 and 0.1 s intervals, while the second set used 50 ms flashes delivered in the same manner. (**B**) Two representative ERG traces evoked by 500 ms flashes. The off-transients are enlarged to the right. The lower trace displays light-off oscillations (arrows), while the other does not. (**C**) Two representative traces evoked by 50 ms flashes (lower panel). The on-transients are enlarged to the right. The trace above shows light-on oscillations (arrows), while the trace below does not.

A second form of oscillation was observed during a sequence of short (50 ms) light flash trains (inter-flash interval: 2.5, 0.5 and 0.1 s; Figure 3A). During these short flashes full ERG waveforms were not evoked; instead, only the on-transient and the initial portion of the RP were observed (Figure 3C). Successively evoked on-transients sometimes displayed a high frequency oscillation (between 100-200 Hz) immediately after the peak of the on-transient (Figure 3C). These “light-on” oscillations were frequently observed between the 2^nd^ and 5^th^ light flash of the 10 flash train, and were approximately 0.3 mV in size, smaller than the light-off oscillations. Light-on oscillations tended to appear in the later stimulus trains (0.1 s inter-flash intervals, Figure 3A). However, the occurrence of light-on oscillations was not necessarily dependent on the inter-flash interval. When the stimulus protocol was reversed, the oscillations were observed during 2.5 s intervals, although the rate of observation was reduced.

The novel ERG oscillations evoked by repetitive light flashes we observed prompted us to eliminate a number of artefactual possibilities that could potentially drive the phenomena (data not shown): 1) <u>The flash intensity oscillated</u>. Using a linear phototransistor circuit (Panasonic AMS302) to measure light output, we noted that the flash intensity was roughly square, with a slight (~5%) overshoot during the initial phase. The light output never oscillated during periods of observed ERG oscillation. 2) <u>Oscillation was specific to a single electrophysiological rig</u>. Both types of ERG oscillations were observed on each of the three rigs used in the *Neurobiology Laboratory* class. Furthermore, using the same light flash protocols, ERG oscillations were observed using an electrophysiology rig in Dr. Chun-Fang Wu’s laboratory. 3) ERG oscillations require a specific light source. Reducing light intensity by 50% using a neutral density (ND) filters did not disrupt the ERG oscillations. Furthermore, oscillations could be triggered by a wide range of light wavelengths. Both a blue LED (wavelength: 470 ± 20 nm, Phillips Luxeon Rebel LED), as well as a white LED passed through a long-pass filter (Kodak Wratten #12, cutoff wavelength ~510 nm) were able to trigger both types of oscillations. Indeed, in our experiences, across an assortment of “white” LEDs, all were able to trigger ERG oscillations in WT flies during repetitive light flashes.

Given the frequent observation of both light off- and light on-oscillations in the WT strain *Canton-S*, we wanted to determine whether these oscillations could be observed in other *Drosophila* WT strains. We examined ERG waveforms of two other commonly used laboratory *D. melanogaster* strains: *Oregon-R* and *Berlin* using the repetitive light flash protocol described above. Importantly, we observed both light off- and light on-oscillations were in these strains, and the frequencies of both types of oscillation were within the range seen in the *Canton-S* strain (Table 1). These findings suggest that the occurrence of either form of oscillation is likely not due to a particular genetic background in *D. melanogaster*. We then extended our analysis of ERG oscillations to include WT strains of four additional *Drosophila* species *euronotus, funebris, americana*, and *hydei*. As listed in Table 1, all four species displayed both kinds of ERG oscillations. However, in several cases the observed oscillations varied in terms of frequency compared to *melanogaster* (Table 1). Although additional work is required to systematically characterize the mechanisms for this variation, it is clear from our initial results that ERG oscillations triggered by repetitive stimulation may be widely observed across *Drosophila* species.

**Table 1.**

| *Species (Strain)* | **Light-off oscillation** |  | **Light-on oscillation** |  |
| --- | --- | --- | --- | --- |
|  | **Fraction** | **Freq. Range (Hz)** | **Fraction** | **Freq. Range (Hz)** |
| *D. melanogaster (Canton-S)* | 19/20 | 70-110 | 9/19 | 100-200 |
| *D. melanogaster (Oregon-R)* | 4/4 | 80-100 | 2/4 | ~140 |
| *D. melanogaster (Berlin)* | 5/5 | 70-100 | 3/5 | 100-150 |
| *D. euronotus* | 4/5 | 60-100 | 5/5 | 50-150* |
| *D. americana* | 5/5 | 70-150 | 5/5 | 60-150* |
| *D. hydei* | 5/5 | 80-120 | 5/5 | 100-120 |
| *D. funebris* | 4/5 | 80-120 | 5/5 | 60-120 |

\* Substantial frequency variation within individuals

### Enhancement of light-off oscillations and suppression of light-on oscillations in the ERG mutant rosA

Oscillations within ERG waveforms are a well-described phenotype of a number of visual mutants (Kelly & Suzuki, 1974; Wu & Wong, 1977; Leung, Geng, & Pak., 2000). Perhaps the best characterized of these are mutants of the gene *rosA* (Wu & Wong, 1977; independently isolated as *inebriated*, Stern & Ganetzky 1992) which encodes a putative Na+/Cl'-dependent solute transporter (Soehenge *et al*., 1996; Burg *et al*., 1996) potentially involved in neurotransmitter transport (Huang *et al*., 2002). We wanted to compare the ERG waveform oscillations in *rosA* mutants with the light-on and light-off oscillations in WT flies described here. Consistent with previous reports (Wu & Wong, 1977; Gavin, Arruda, & Dolph, 2007), our ERG recordings of the allele *rosA*^*p*213^ revealed that 500 ms light flashes did not evoke on- or off-transients, but recruited a prominent oscillation during the RP phase of the waveform (~ 3 mV, Figure 4A). The frequency of the RP oscillation varied somewhat between individuals (40 to 90 Hz), but ceased within 10 ms of the end of the light flash. Using the repetitive light flash protocols previously described (Figure 3A), we found several differences in how light-off and light-on oscillations were recruited in *rosA*. Specifically, the repetitive 500 ms flash protocols (inter-flash interval: 2.5 or 0.5 s) were able to recruit strong light-off oscillations during the repolarization phase of the waveform, often lasting tens of milliseconds (Figure 4A-B). Within the same individual, the frequency of the light-off oscillations was lower than corresponding RP oscillation frequency (Figure 4B). Compared to WT counterparts, the light-off oscillations in *rosA* were significantly larger in amplitude (~3 mV vs < 1 mV) and generally had a lower oscillation frequency (~50 Hz, Figure 4B). In contrast to the enhanced light-off oscillations in *rosA*, we found that the 50 ms light flash protocols could not recruit light-on oscillations in *rosA*. Taken together, these results seem to indicate a critical role for the *rosA* gene product in shaping light-off and light-on oscillations in addition to suppressing RP oscillations in WT ERG waveforms.

**Figure 4.**
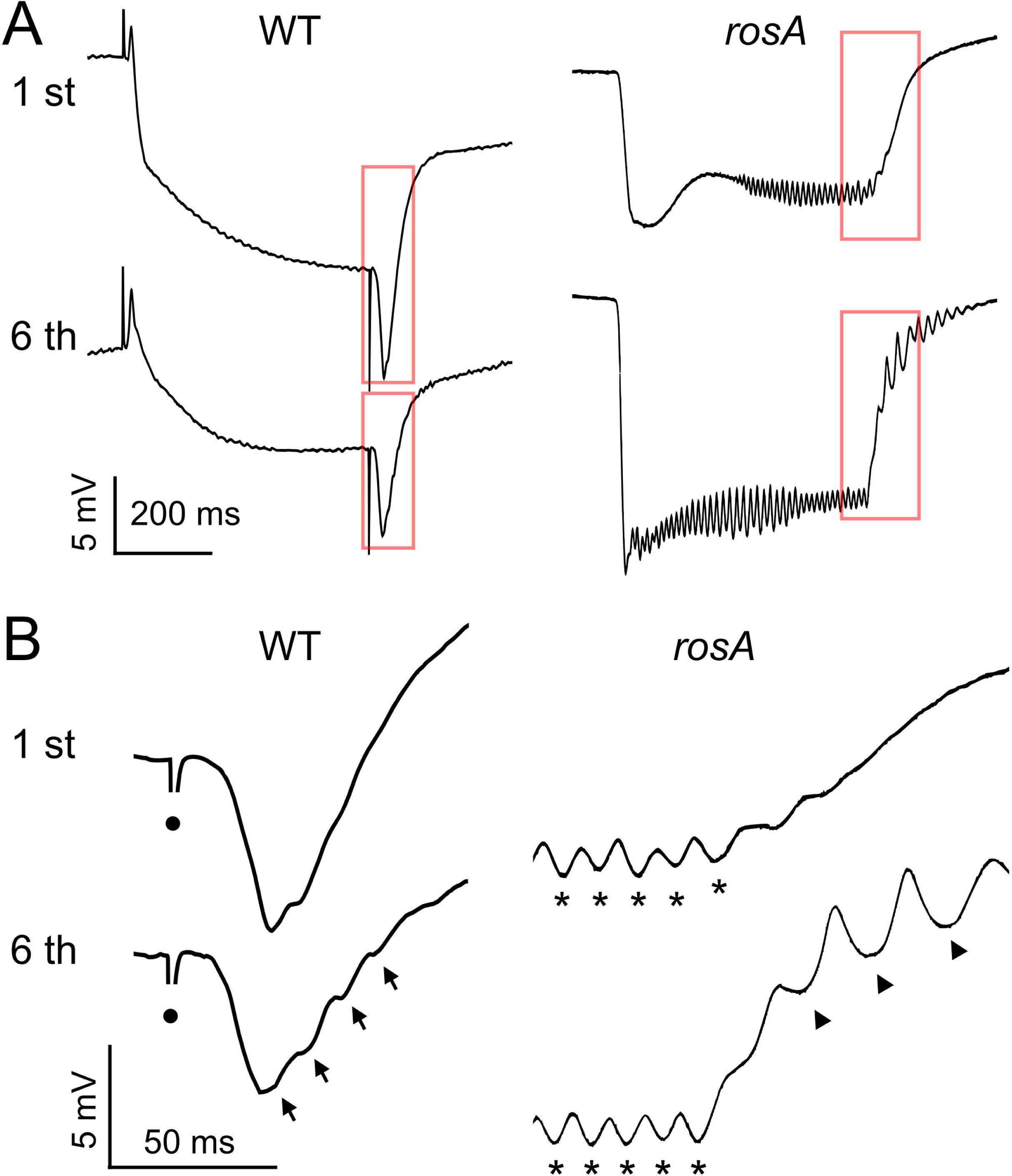
Light-off oscillations in the visual system mutant *rosA*. (**A**) ERG waveforms evoked by a 500 ms light flash in WT and *rosA* mutants. Upper traces: 1^st^ flash, lower traces: 6^th^ flash; inter-flash interval: 500 ms. Note the oscillations during the RP and absence of transients in *rosA*. (**B**) Expansion of the off-transient and repolarization. Arrows indicate light-off oscillations, asterisks indicate RP oscillations in *rosA*, and circles indicate light-off artifact.

### Age-dependent suppression of ERG waveform oscillations

Although repetitive light flashes evoke light-off oscillations reliably and often trigger light-on oscillations in young flies, we noticed that these oscillations were largely absent from the older WT flies we examined. As shown in Figure 5A, in 29 °C-reared WT flies, we observed light-off oscillations in 23 out of 24 flies less than 15 d old (corresponding to ~50% of the median lifespan), while in older flies, only 3 out of 11 displayed oscillation. Interestingly, the frequency of light-off oscillation was also age dependent, with relatively younger flies displaying higher frequency oscillations compared to older counterparts (Spearman’s rank correlation test, *p* < 0.05). The proportion of flies displaying light-on oscillations also decreased with age: 10 out of 23 flies for flies less than 15 d old vs. 1 out of 9 for flies older than 15 d. However, we did not detect a significant age dependence in the light-on oscillation frequency (Figure 5B). Based on these observations, loss of light-on or light-off transients may serve as a marker of age-dependent alterations in the *Drosophila* visual system alongside the attenuation of ERG transients.

**Figure 5.**
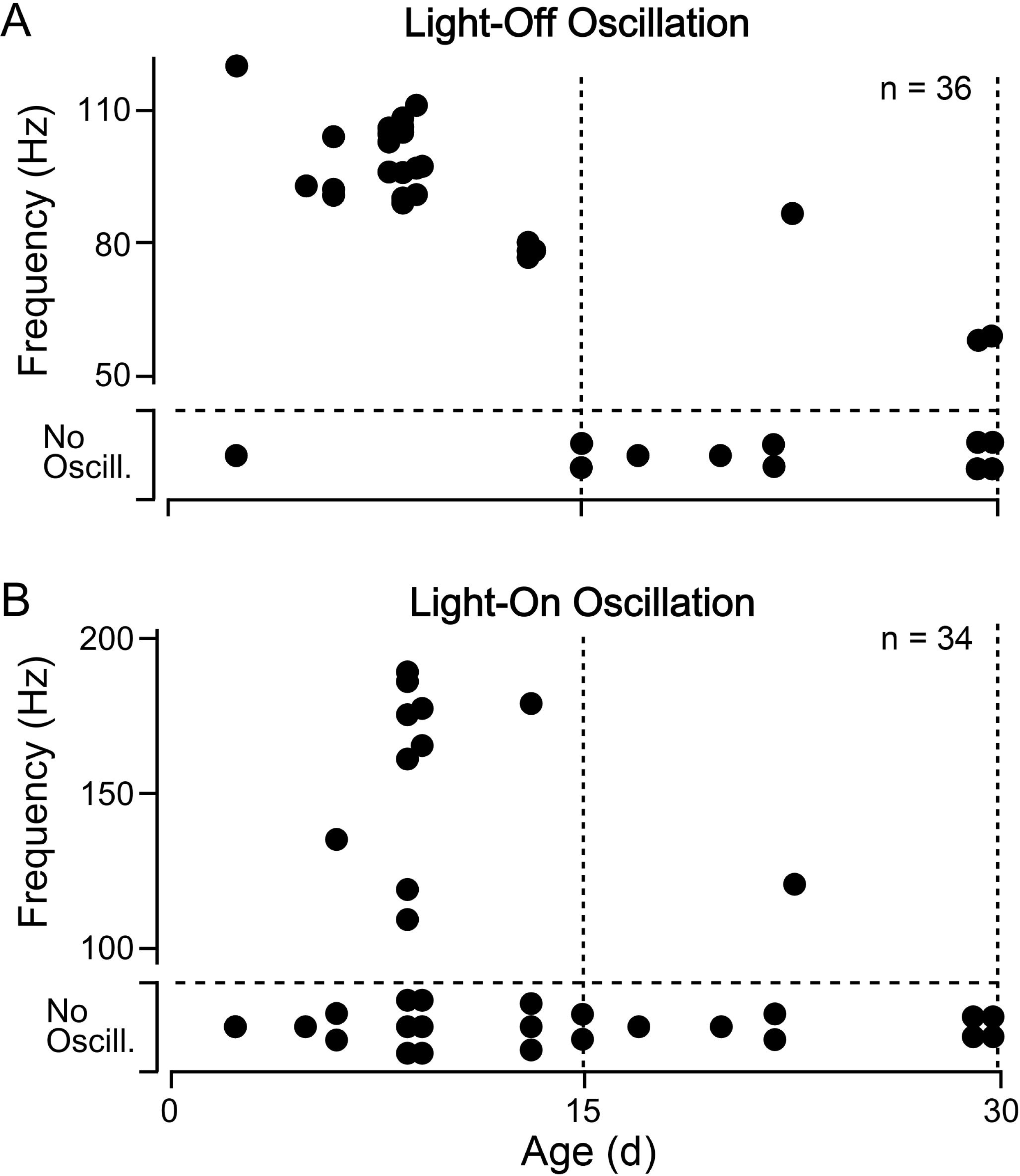
Age-related changes in light-off and light-on oscillations (**A**) Scatterplot of age versus oscillation frequency for light-off oscillations in WT flies and (**B**) light-on oscillations. Sample sizes as indicated in upper-right corner. Note that flies that did not display oscillations are indicated at the bottom of each plot. A Fisher’s exact test revealed that both light-on and light-off oscillations were more frequently observed in younger flies (p < 0.001 for light-off oscillation, *p* < 0.05 for light-on oscillation).

## Discussion

Our study compares the progression of age-related changes in phototactic behavior with changes in the *Drosophila* ERG waveform. We observed an age-related decline in phototaxis during which negative geotaxis within the same population was consistently stronger (Figure 1). The relative strength of negative geotaxis suggests that the decline in phototaxis may be due to disruption of requisite sensory inputs rather than motor system function. Indeed, previous findings using different methodologies have shown a similar decline in phototaxis which precedes a loss of motor coordination in aging flies (Leffelaar & Grigliatti, 1983; Arking & Wells, 1990). Interestingly, we noted that the decline in phototaxis was paralleled with the age-related decrease of on- and off-transients in ERG waveforms (Figure 2). On- and off-transients serve to indicate integrity of synaptic transmission between photoreceptors and 2^nd^ order targets (Coombe, 1986), and disruption of transients leads to distinct phototaxis deficits (Pak, 1975). These results suggest that the loss of ERG transients may contribute to the observed loss of phototaxis. Among the 23 °C-reared WT population aged beyond 40 d, we found that flies with a phototaxis score of ‘3’ had larger average off-transient amplitudes than those with a score of ‘0’. However, a limited sample size of old flies displaying good phototaxis precluded robust statistical analysis, and we identified a few individuals with a ‘poor’ phototaxis score which displayed robust transients. Further study, perhaps utilizing a behavioral assay sensitive to individuals such as the opto-motor response in a tethered flight simulator (Götz, 1968; Götz, Hengstenberg, & Biesinger, 1979) may more directly correlate the loss of transients with visually mediated behaviors within individual aged flies.

Notably, a loss of ERG transients coupled with a decline in visually mediated behaviors is a phenotype associated with a number of neurodegeneration mutants (e.g. *optomotor blind*, Heisenberg, Wonneberger, & Wolf, 1978; *swisscheese*, Kretzschmar, Hasan, Sharma, Heisenberg, & Benzer, 1997) which also display clear disruptions in photoreceptor or laminar structure (Pflugfelder *et al*., 1990; Pflugfelder *et al*., 1992; Kretzschmar, *et al*., 1997; Kretzschmar 2009). It is possible that similar structural defects in photoreceptor, laminar and/or surrounding glial cells may contribute the observed decline in transients in aged flies. Furthermore, direct monitoring of the post-synaptic potentials in laminar and/or medullar cells perhaps via direct patch recordings (Skingsley, Laughlin, & Hardie, 1995; Tuthill, Nern, Rubin, & Reiser, 2014) or Ca^2+^ imaging (Clark, Bursztyn, Horowitz, Schnitzer, & Clandinin, 2011) would be potentially useful in pinpointing the physiological basis for this decline. Previous work in other systems have shown age dependent changes in field recordings of synaptic transmission (Koss, Drever, Stoppelkamp, Riedel, & Platt, 2013; Piskorowski *et al*., 2016) and circuit activity (Hughes & Cayaffa, 1977; Landholt & Borbely, 2001).

Our repetitive light flash protocol (Figure 3A) was inspired by previous studies that used repetitive stimulation to uncover age- and activity-dependent alterations in synaptic physiology including the crayfish NMJ (Govind, 1992; Atwood, 1992), along the *Drosophila* giant fiber jump and flight escape circuit (Martinez *et al*., 2007; Ruan, 2008) and motor circuits recruited during seizure discharges (A.I., unpublished observations). Significantly, the repetitive light flash protocol resulted in our discovery of two novel ERG oscillations in WT flies (Figure 3B). Light-off oscillations, the more prominent of the two, appeared during trains of 500 ms flashes, with amplitudes generally growing during each train (up to 1 mV). Light-on oscillations, in contrast, were often observed during the 2^nd^ -5^th^ of the short duration light flashes (50 ms). The light flash history-dependence of these oscillations suggests that one of the conditions to induce ERG oscillations is repeated activity at the retina.

Although oscillations can be observed in the electrical activity of isolated mutant photoreceptor cells with the use of intracellular recordings (Wu & Wong, 1977), the ERG waveform oscillations likely reflect collectively synchronous activity of many photoreceptor cells and their synaptic connections. The relative resistance between photoreceptor cells is much smaller than the resistance across the basement membrane. Thus, when the probe electrode and the ground electrode are placed on opposite sides of the retina’s basement membrane, the resulting ERG waveform represents the combined activity of photoreceptors and their synaptic targets (Heisenberg, 1971; Stark & Wasserman, 1974). Oscillations may arise through an interaction between photoreceptor activities, such as feedback amplification, which serves to synchronize them. A glial cell syncytium connected through gap junctions (Stebbings *et al.*, 2002; Chaturvedi, Reddig, & Li, 2014) could contribute an important mechanism to synchronize photoreceptor and/or laminar activity, facilitating the reported ERG oscillations.

The two separate ERG oscillations identified may involve distinct combinations of physiological mechanisms involving photoreceptor cells, postsynaptic monopolar laminar cells, as well as surrounding glial cells (Heisenberg, 1971; Coombe, 1986; Xiong & Montell, 1995). The different stimulus recruitment protocols and oscillation frequencies of light-on and light-off oscillations (Figure 3, Figure 5) suggest that they arise through independent mechanisms. Indeed, we noted several WT individuals which displayed light-off but not light-on oscillations (Table 1). As mentioned above, light-on oscillations were absent in flies with attenuated on-transients including both *rosA* mutants and older WT flies, suggesting that light-on oscillations require intact on-transients. (Preliminary analysis of the transient-less mutant lines *hdc*^*p*211^ and *ort*^*k*84^ also indicates a loss of light-on oscillations; I.M., unpublished observations.) Light-off oscillations, in contrast, do not require intact off-transients, as *rosA* mutants which lacked off-transients displayed strong light-off oscillations in response to repetitive light flashes (Figure 4). However, our observations indicate a potential role for the *rosA* gene product in the light-off oscillations. In *rosA* mutants, the light-off oscillations are distinct from previously described RP oscillations in terms of frequency and location within the ERG waveform (Figure 4). Furthermore, the frequency of light-off oscillations in *rosA* is reduced and the amplitude is increased compared to WT light-off oscillations. Although the precise physiological mechanisms by which *rosA* mutations induce RP oscillations remain unclear (Gavin *et al*., 2007), the *rosA* gene product along with other neurotransmitter transporters (e.g. *carT*, see Xu *et al*, 2015; Chaturvedi *et al*., 2016) may play an important role in shuttling histamine and related metabolites across the plasma membranes of photoreceptors and surrounding glia in the visual system—a potentially important mechanism in generating ERG waveform oscillations.

## Acknowledgments

We thank Prof. Chun-Fang Wu for supervising the *Neurobiology Laboratory* course and for providing his support and encouragement to develop this work. We also thank Prof. Bernd Fritzsch, Prof. Michael Dailey and Dr. Olga Miakotina for organizational and technical assistance. We thank other participants of the course for their assistance in data collection. AU and AI were supported by the NIH grant AG051513 to Chun-Fang Wu.

---

**Supplemental Figure 1** Age-dependent alterations of 23 °C-reared WT ERG waveforms. The amplitudes of (**A**) RP, (**B**) on-transient, and (**C**) off-transient. Trend-lines indicate a Gaussian-kernel running average (σ = 5 d). n = 19 flies.

